# Independent avian epigenetic clocks for aging and development

**DOI:** 10.1101/2024.09.20.614163

**Authors:** Ayke Haller, Judith Risse, Bernice Sepers, Kees van Oers

## Abstract

Information on individual age is a fundamental aspect in many ecological and evolutionary studies. However, accurate and non-lethal methods that can be applied to estimate the age of wild animals are often absent. Furthermore, since the process of ageing is accompanied by a physical decline and the deterioration of biological functions, the biological age often deviates from the chronological age. Epigenetic marks are widely suggested to be associated with this age-related physical decline, and especially changes in DNA methylation are suggested to be reliable age-predictive biomarkers. Here, we developed separate epigenetic clocks for ageing for development in a small passerine bird, the great tit (*Parus major*). The ageing clock was constructed and evaluated using erythrocyte DNA methylation data of 122 post-fledging individuals, and the developmental clock using 67 pre-fledging individuals from a wild population. Using a leave-one-out cross validation approach, we were able to accurately predict the ages of individuals with mediation absolute deviations of 0.40 years for the ageing and 1.06 days for the development clock. Moreover, using existing data from a brood-size manipulation we show that nestlings from reduced broods are estimated to be biologically older compared to control nestlings, while they are expected to have higher fitness. These epigenetic clocks provide further evidence that, as observed in mammals, changes in DNA methylation of certain CpG sites are highly correlated with chronological age in birds and open up new avenues for broad applications in behavioural and evolutionary ecology.

## 1. Introduction

Age is a fundamental aspect in the field of ecology and evolution and biological characteristics, such as size and sexual maturity, are directly linked to an organisms’ age (Stearns, 1976). However, these characteristics change during an organisms’ life, affecting the survival and reproductive effectiveness of an individual and hence the population structure as a whole (Heydenrych et al., 2021). Information of chronological ages can thus be important to determine the overall population viability and the mortality of individuals. However, accurate estimations of chronological ages in wild populations can only be derived from long-term tracking studies, in which individuals have been marked at birth. Alternative methods estimate ages in wild animals, such as teeth measurements in deer (Pérez-Barbería et al., 2014) and foxes (Chevallier et al., 2017), often involve high uncertainties and can require invasive capture and release strategies (Kukalová et al., 2013), while other methods necessitate lethal sampling (Gunn et al., 2008; Hammerschlag & Sulikowski, 2011). The absence of accurate and non-lethal methods to determine the chronological age, necessitates the development of novel tools that can be applied to wild populations.

Because the process of ageing, which is defined by a physical decline that results in the deterioration of biological functions (often referred to as senescence), the cellular state may vary between individuals of the same chronological age (Kirkwood, 2005). Moreover, the rate of senescence can also vary between individuals within a population (Seehuus et al., 2006; Lowsky et al., 2014). This emphasises that chronological age is an imperfect substitute to measure age and the process of aging (Horvath & Raj, 2018). The biological or cellular state of an individual, here referred to as biological age, might therefore act as a better predictor for an individuals’ health and survival. To quantify biological age, specific sets of age-related biomarkers have to be identified (Baker and Sprott, 1988). In addition, an important distinction needs to be made between the age-related changes during the developmental stages of an individual and the overall biological deterioration associated with ageing. Age-related changes in biological processes during development might, in contrast to changes during senescence, have a positive influence on the cellular state and fitness. This either because of increased sizes after development, or the risks involved with prolonged growing periods (Gebhardt-Henrich & Richner 1998). Therefore, the distinction needs to be made between biomarkers identified during these different life stages, which, to the best of our knowledge, have not been done in one system.

Biochemical mechanisms, particularly some epigenetic marks, are widely suggested to be associated with an organisms chronological age (Oberdoerffer & Sinclair, 2007; Campisi & Vijg, 2009). Epigenetic mechanisms refer to those mechanisms that can modify the expression of genes by either silencing transcription or translation, without changing the nucleotide sequence of a genome (Jablonka & Lamb, 1998). Epigenetic modifications include, for instance, histone modifications, non-coding RNA, and DNA methylation (DNAm), the latter of which being the most studied modification that is found in organisms across various taxa (Bellizzi et al., 2019). DNAm is the addition of a methyl group (-CH3) to a DNA nucleotide, either a cytosine or an adenine (Jarman et al., 2015). In vertebrates, DNAm mostly occurs when a cytosine (C) is followed by a guanine (G), separated by a phosphate (p) group, which is commonly referred to as a CpG site (hereafter CpGs). In recent years an increasing number of studies have demonstrated that some CpGs can be subjected to an age-related decrease (hypomethylation) or increase (hypermethylation) in DNAm. This highlights the potential of DNAm as a promising biomarker to investigate biological age (Horvath, 2013). Epigenetic drift is the age-associated alteration of DNAm (Vaiserman, 2018), which is often studied in relation to the patterns at a large number of CpGs distributed throughout the whole genome (e.g., Gryzinska et al., 2013; Shimoda et al., 2014). However, specific CpGs that correlate with chronological age do not necessarily exhibit a similar trend in the age-associated alterations (Horvath, 2013), and this distinction cannot be accounted for in an analysis of the global patterns alone.

Clock type biomarkers, or epigenetic clocks, are an example of specific sets of CpGs from which the DNAm levels collectively exhibit a high correlation with chronological age. Compared to other potential estimators for biological age, including telomere length and metabolomics-based predictors, epigenetic clocks demonstrated to be the most promising avenue as a molecular estimator of biological age (Jylhävä et al., 2017), and DNAm as the most reliable age-predictive biomarker (Lee et al., 2016). This was demonstrated by the introduction of epigenetic clocks for humans, and since then there has been a growing interest in the development of these clocks for other species as well. To this regard, similar clock models have been developed for model species such as mice (Petkovich et al., 2017; Wang et al., 2017) and rats (Levine et al., 2020), and several have already been developed for wild animals, however, predominantly for mammalian species (e.g., see Anderson et al., 2021; Pinho et al., 2022; Peters et al., 2023).

In this study we developed two distinct epigenetic clock models using DNAm patterns from blood samples of the great tit (*Parus major*), a passerine bird, by applying a penalised regression to identify the most informative CpGs. Bird systems are ideal models to study the disparity between the rate of ageing during the developmental phases and throughout the whole life, since they develop outside of the mother and therefore epigenetic information can be retrieved for both distinct post-natal phases. We produced a high-quality developmental epigenetic clock using 67 pre-fledging individuals of different ages (in days) and an ageing epigenetic clock for 122 post-fledging individuals (in years), originating from a long-term nest box population in the Netherlands, using a cost-effective reduced representation bisulfite method (EpiGBS2, Gawehns et al., 2022). We then applied the developmental clock by using samples from nestlings that were part of a brood size manipulation experiment (Sepers et al., 2023a & 2023b) to test the hypothesis that a higher developmental epigenetic age is associated with lower nutritional stress during early rearing.

## 2. Material and Methods

### 2.1 Samples, DNA extraction and sequencing

#### 2.1.1 Sample collection

Individuals included in this study were part of a long-term nest box population of great tits (*P. major*) in the Westerheide estate near Arnhem, the Netherlands (52°01′00N, 05°50′30E). We used blood samples of 134 post-fledging individuals that were collected in 2018. The chronological ages of these individuals ranged between 0.13 and 6.03 years (49 and 2200 days). 62 of these individuals were parents of broods that were used in a brood size manipulation as described in detail in Sepers et al. (2023a). In short, in a partial cross-foster design nestlings of the same hatching date and similar brood sizes were assigned to a cross-foster pair one or two days after hatching. Nestlings were partially cross-fostered and brood sizes were manipulated by enlarging the brood size by three nestlings, reducing the brood size by three nestlings or by retaining the original number in the brood. The parents were caught using spring traps placed in the next boxes, after which two blood samples of approximately 10µL each were taken from the brachial vein, as described in detail in Sepers et al. (2023a). The remaining 72 individuals were their fledged offspring originating from the same brood size manipulation experiment. Several weeks after fledging, these individuals were recaptured using mist nets, and a blood sample of approximately 20µL was taken from the brachial vein, as described in detail in Sepers et al. (2024a). For 96 post-fledging individuals, the precise hatching date was known, allowing for the accurate determination of their age at the moment of sampling. For the remaining 38 individuals, the precise hatching date was not registered, however, the year in which these individuals hatched was known based on their ring numbers or moulting pattern. Therefore, we used the average hatching date of the year that these individuals hatched to calculate their age at the moment of sampling.

Furthermore, we used blood samples from nestlings (*n* = 618) during several experimental studies conducted throughout 2018, 2019 and 2020. In 2018, blood samples of approximately 10µL were taken from 222 nestlings from the brachial vein at 14 (*n* = 211) or 15 (*n* = 11) days after hatching, as part of the previously described brood size manipulation experiment and stored in cell lysis buffer. Furthermore, in 2019, an experiment was conducted investigating the effect of food deprivation (Sepers unp.). For that, nestlings were assigned to a food deprivation treatment (*n* = 108) or control group (*n* = 108). The individuals from the food deprivation treatment were denied food by placing a metal cage on top of them during the 7^th^, 8^th^ and 9^th^ day after hatching for a period of two, two and a half, and three hours, respectively, allowing the parents to feed the other half of the brood. Blood samples of approximately 10µL were taken from the brachial vein from all individuals 7, 9 and 14 days after hatching and stored in cell lysis buffers. Finally, in 2020, an experiment was conducted to investigate the effects of testosterone manipulation of egg yolk androgen levels. Eggs were subjected to experimental manipulation with either 5µL sesame oil (control; *n* = 90) or a testosterone solution (*n* = 90) injected using a 10µL syringe with a 26G needle (Sepers et al. 2024b). Subsequently, blood samples of approximately 10µL were taken from individuals six (*n* = 140) or seven days (*n* = 40) after hatching. For all pre-fledglings the precise date of hatching was known, which allowed us to accurately determine their chronological age at the moment of sampling which ranged between 6 and 15 days.

#### 2.1.2 DNA extraction and sequencing

Library preparation was conducted following the laboratory protocol described in Gawehns et al. (2022) and Sepers et al. (2023b). Briefly, FavorPrepTM 96-Well Genomic DNA Extraction Kit (Favorgen) was used to isolate DNA from each blood sample. 800ng DNA was extracted and digested using the restriction enzyme MspI (NEB) that cuts DNA at C^CGG motif. Beads (0.8× AMPure XP beads; Beckman Coulter) were used to remove large fragments, which were then ligated to customised barcoded adapters. For each library, samples were pooled (multiplexed), and small fragments (<60bp) were removed with Nucleospin Gel and PCR kit (Macherey-Nagel). Next, the nick between 5’ ends of the adapter and 3’ ends of the fragments were repaired, and treated with sodium bisulfite. KAPA HIFI Uracil + hotstart ready mix (Roche) was used to amplify the libraries after the sodium bisulfite conversion during 15 PCR cycles. Then the libraries were cleaned with NucleoSpin Gel and PCR Cleanup Kit and 0.8x AMPure XP beads. Finally, the quantity and quality of each library was determined with qPCR quantification (KAPA Library Quantification Kits, Roche).

For samples collected in 2018, the bioinformatic analysis was performed as described in Sepers et al. (2023a, 2023b & 2024a). The raw reads were demultiplexed, checked for quality and adapter content, and merged using epiGBS2 following the details described *for P. major* samples in Gawehns et al. (2022). Subsequently, short reads (< 20bp) were removed, and the 3’ end adapter and Illumina sequence were trimmed using Cutadapt v2.10 (Martin, 2011). An additional 10bp were then removed from the 3’ end. Quality of the raw and cleaned reads were assessed with FastQC v0.11.8 (Andrews 2010), FastQ screen v.0.11.1 in bisulfite mode (Wingett and Andrews 2018), and MultiQC v1.8 (Ewels et al., 2016). Next, the reads were aligned in paired-end and non-directional mode to the *P. major* reference genome v.1.1 (GCF_001522545.3; Laine et al., 2016) using Bismark v0.22.3 (Krueger & Andrews, 2011), and methylation was exclusively called in CpG context while removing the overlap of the reads and ignoring the first four bp’s. For the samples collected in 2019 and 2020, the raw reads were demultiplexed, checked, trimmed, filtered, merged, aligned, and called for methylation using the reference branch of the bioinformatic pipeline epiGBS2 (Gawehns et al., 2022), using the custom settings to call methylation exclusively in CpG context while removing the overlap of the reads and ignoring the first four bp’s.

### 2.2 Sample selection and filtering methylation calls

EpiGBS2 is a reduced representation DNAm analysis tool that prioritises efficiency and cost-effectiveness in library preparation and is known to induce a considerable between-sample variation in the uniformity of the CpGs that are called. However, for inclusion in the clock model analysis, it is essential that CpGs are covered throughout all samples. We therefore made different selections to filter the methylation calls of post- and pre-fledging samples, to maximise the number of CpGs that could be used to train the clock models.

#### 2.2.1 post-fledging

The R package *methylKit* v.1.28 (Akalin et al. 2012) was used to merge complementary CpG dinucleotides. Next, using a custom R script, we removed each CpG with a coverage of less than 10 and with a coverage over the 99.9 percentile. For each CpG, we calculated the methylation ratio by dividing the number of methylated Cs by the total coverage of that site. Finally, to exclude sites with a low variation in methylation, which is required to find correlations with age, we removed sites with a mean methylation level lower than 5%, or higher than 95% and sites with a standard deviation of less than 0.05.

#### 2.2.2 pre-fledging

Unlike the post-fledging samples, here we first discarded CpGs with a coverage of less than 10 and with a coverage over the 99.9 percentile, before merging complementary CpG dinucleotides. For each CpG, we calculated the methylation ratio by dividing the number methylated Cs by the total coverage of that site. However, even after these filtering decisions it proved difficult to find uniform sites covered throughout all samples. We therefore randomly divided the samples into four groups stratified for age, using the R package *caret* v.6.0-94 (Kuhn, 2008), and evaluated the number of sites that were uniform within each subset. This process was repeated for 500 iterations to increase the probability of finding an optimal subset that contained a high number of CpGs. Finally, to exclude sites with low variation in methylation, we removed sites with a mean methylation level lower than 5% or higher than 95% and with a standard deviation of less than 0.05.

### 2.3 Epigenetic clocks and age estimations

We constructed two epigenetic clocks, an ageing clock using the age in years of post-fledging individuals, and a development clock, using the age in days of pre-fledging individuals. The clock models were constructed in R software using penalised regression implemented with the R package *glmnet* v.4.1-8 (Friedman et al. 2010). This technique allows the user to manually tune the two hyper-parameters alpha (α) and lambda (λ). The α parameter controls the mixing between two types of regression techniques, i.e., ridge regression (α = 0) and least absolute shrinkage and selection operator (LASSO; α = 1). Ridge regression results in models that retain all predictor variables but shrinks correlated variables towards each other, while LASSO generally shrinks coefficients of correlated predictors towards zero, resulting in simpler models that retain fewer variables. The mixing parameter used in both regression techniques can be combined to create a more regularised model, referred to as an elastic net model which provides the flexibility to set the α between 0 - 1. Before constructing the final clock models, we investigated the effect of different α values between 0 – 1 on the model performance, by running a model with all samples, for the ageing and development clock separately, using the *cv.glmnet* function with internal 10-fold cross-validation for each α in increments of 0.1. This process was repeated for 100 iterations, and we selected the α of 0.2 for both clock models as a compromise between minimising the error margin and retaining CpGs. Furthermore, the λ penalty parameter (lambda.min) was selected from the model that minimalised the mean squared error (MSE). To assess the predictive accuracy of our final clock models, we used these selected parameters in a leave-one-out cross-validation (LOOCV) approach, to predict the epigenetic age of each sample. The accuracies of these predictions were evaluated using the Pearson correlation coefficient (*r*), and the median absolute deviation (MAD) between the predicted epigenetic age and the chronological age. The final versions of the clock models were fitted using the selected hyper-parameters and included all samples.

### 2.4 Statistical analysis

First, we examined whether the retained CpGs of the final development clock model could be detected in samples that were taken from 14-day old individuals. Then to analyse whether experimentally induced nutritional stress during the pre-fledging influenced the epigenetic ages of the individuals out of a control, reduced or enlarged brood size (Sepers et al., 2023a), we used the determined coefficients of the final developmental clock to estimate the epigenetic age of the individuals in which these CpGs were detected. Subsequently, we developed a linear mixed model, using R package *lme4* v.1.1-35.1 (Bates et al. 2015), with the estimated epigenetic age as response variable, treatment as fixed effect (three levels), and the brood of origin as random intercept. This model was then used in a post-hoc analysis, to assess the difference in the estimated marginal means between the three treatment groups using R package *emmeans* v.1.10 (Lenth, 2024), fitting the *p*-values with Fisher’s least significant difference test.

## 3. Results

### 3.1 Universal CpGs

#### 3.1.1 post-fledging

The datafiles of 134 post-fledging samples contained 15,372,018 unique CpGs, with an average of 2,963,751 CpGs per sample (range: 265,600 – 3,840,179 CpGs). In order to maximise the probability of identifying uniform sites throughout the samples, we decided to discard 12 samples that had fewer than 2.5 million CpGs (Figure S1). Discarding samples with a higher number of CpGs caused the loss of a large proportion of older individuals (*n*=122). Merging the complementary CpG dinucleotides resulted in 2,552,100 unique CpGs, from which we only selected the sites that were covered in all 122 samples. After filtering these CpGs based on their coverage (>9X) 35,126 CpGs remained. After excluding the sites with a low variation in methylation ratio, 8,398 unique CpGs remained to construct the ageing clock model (Table S1).

#### 3.1.2 pre-fledging

The datafiles of 618 pre-fledging samples contained 15,372,018 unique CpGs, with an average of 2,784,984 CpGs per sample (range: 808,828 – 3,883,186 CpGs). Here we decided to continue with samples with a minimum of three million CpGs to maximise the probability of obtaining a high number of sites covered in all samples. This left 272 samples with an average of 3,274,660 CpGs, ranging between 3,000,616 – 3,883,186 CpGs per sample for analyses (Figure S2). Merging the complementary CpG dinucleotides resulted in 2,055,751 unique CpGs throughout the 272 samples (Table S2). Randomising these samples into four subsets and evaluating the number of uniform sites during 500 iterations, resulted in four subsets with over 10,000 unique CpGs (Figure S3). From this, we selected the subset that retained the most CpGs, which consisted of 67 samples with 11,116 unique CpGs, and after excluding the sites with a low variation in methylation ratio, 3,184 unique CpGs remained to construct the development clock model (Table S3 & S4).

### 3.2 Epigenetic clock models

The final ageing clock model was constructed using 122 post-fledging samples (58 males and 64 females), ranging in age from less than one year old (minimum age of 0.13 years; 49 days) to six years old (maximum of 6.02 years; 2199 days). This model identified 183 CpGs that correlated with chronological age. The retained CpGs were distributed throughout 28 annotated great tit chromosomes, while 13 CpGs lay on unplaced scaffolds. Age predictions made using LOOCV provided accurate age predictions with a correlation between epigenetic age and chronological age of *r* = 0.84 (95% CI = 0.78 – 0.89, df = 120, p = <0.001) and a MAD of 0.401 years. The model overestimated the epigenetic age of younger individuals of around one year old, while older individuals were largely underestimated compared to their chronological age (Figure 1).

**Figure 1.**
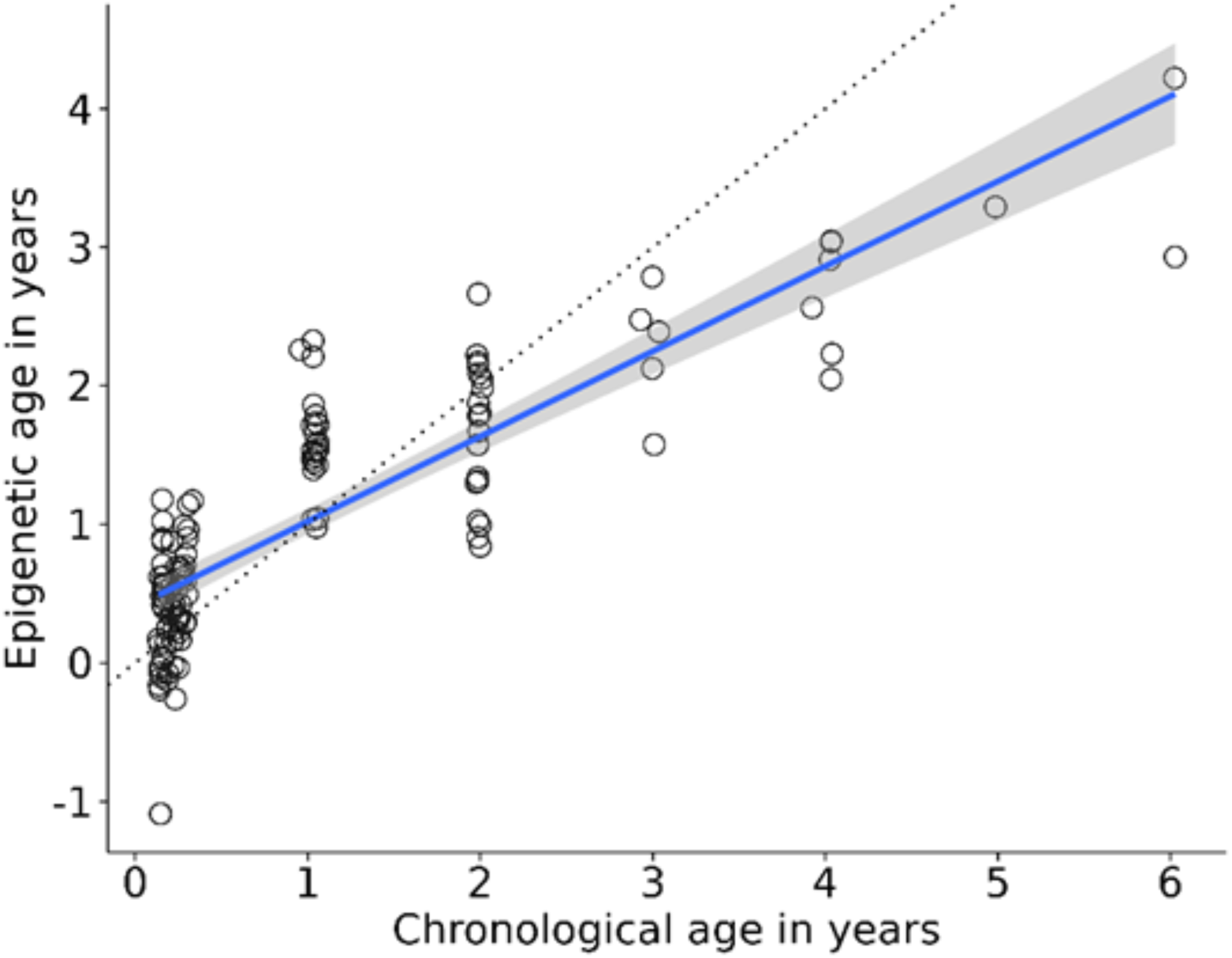
Epigenetic ages of n = 122 post-fledgeling individuals estimated using LOOCV (r = 0.84, MAD = 0.401). Regression line of epigenetic age against chronological age is shown in blue with the standard error in grey. The dotted line represents the regression line if predicted age is identical to chronological age.

The final development clock model was constructed using 67 pre-fledging samples (26 males, 25 females, and 16 of unknown sex) ranging in age from 6 to 15 days. This model identified 115 CpGs that correlated with chronological age. Most of the retained CpGs were distributed throughout 26 annotated great tit chromosomes, while 10 CpGs lay on unplaced scaffolds. Predicted epigenetic ages using LOOCV provided accurate predictions with a correlation between epigenetic age and chronological age of *r* = 0.89 (95% CI = 0.83 – 0.93, df = 65, p = <0.001) and a MAD of 1.061 days (Figure 2). In both the ageing and development clock models, the estimated epigenetic ages showed similar trends of ageing for female and male individuals (Figure 3.)

**Figure 2.**
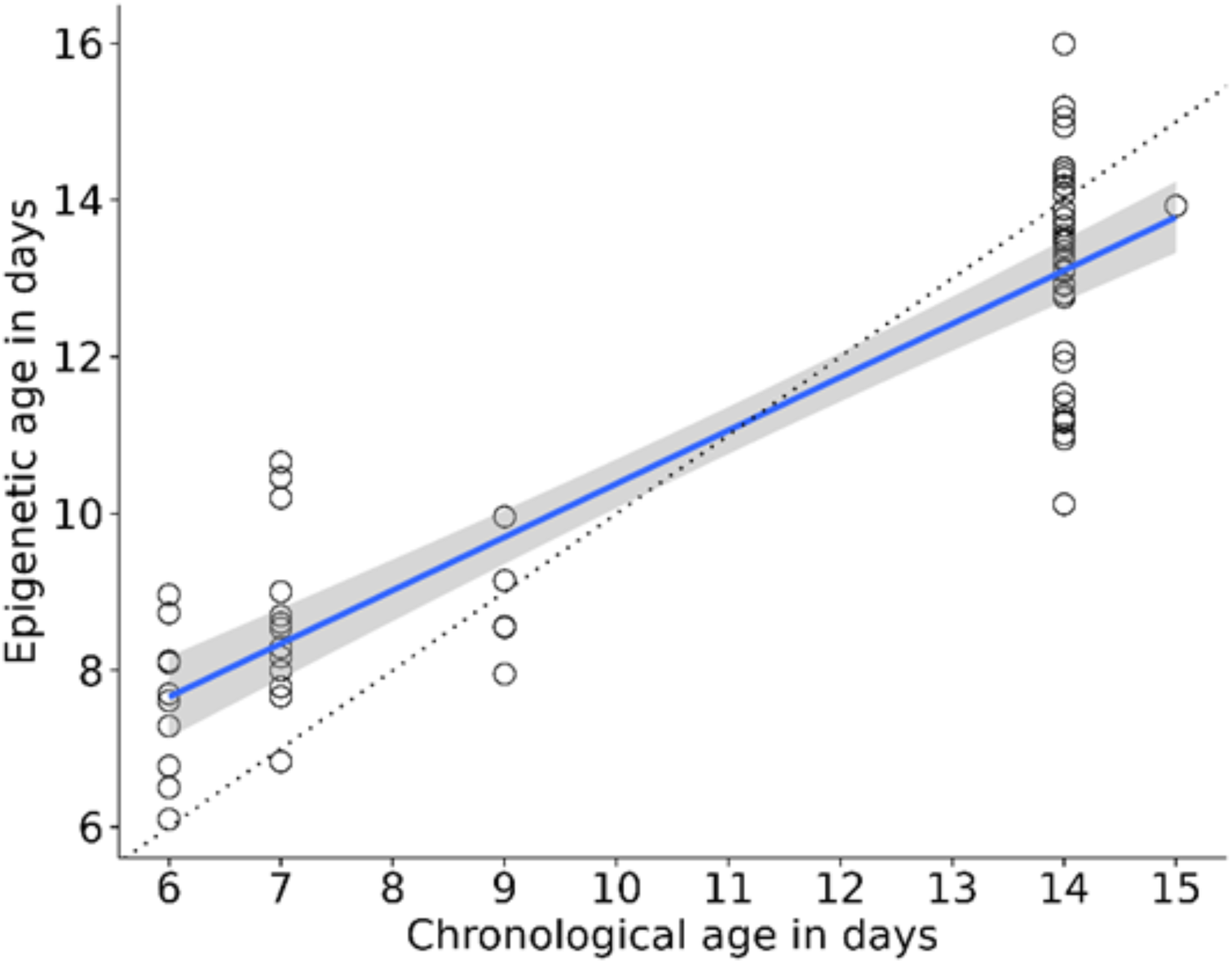
Epigenetic ages of N = 67 pre-fledgeling individuals estimated using LOOCV (r = 89, MAD = 1.06). Regression line of epigenetic age against chronological age is shown in blue with the standard error in grey. The dotted line represents the regression line if predicted age is identical to chronological age.

**Figure 3.**
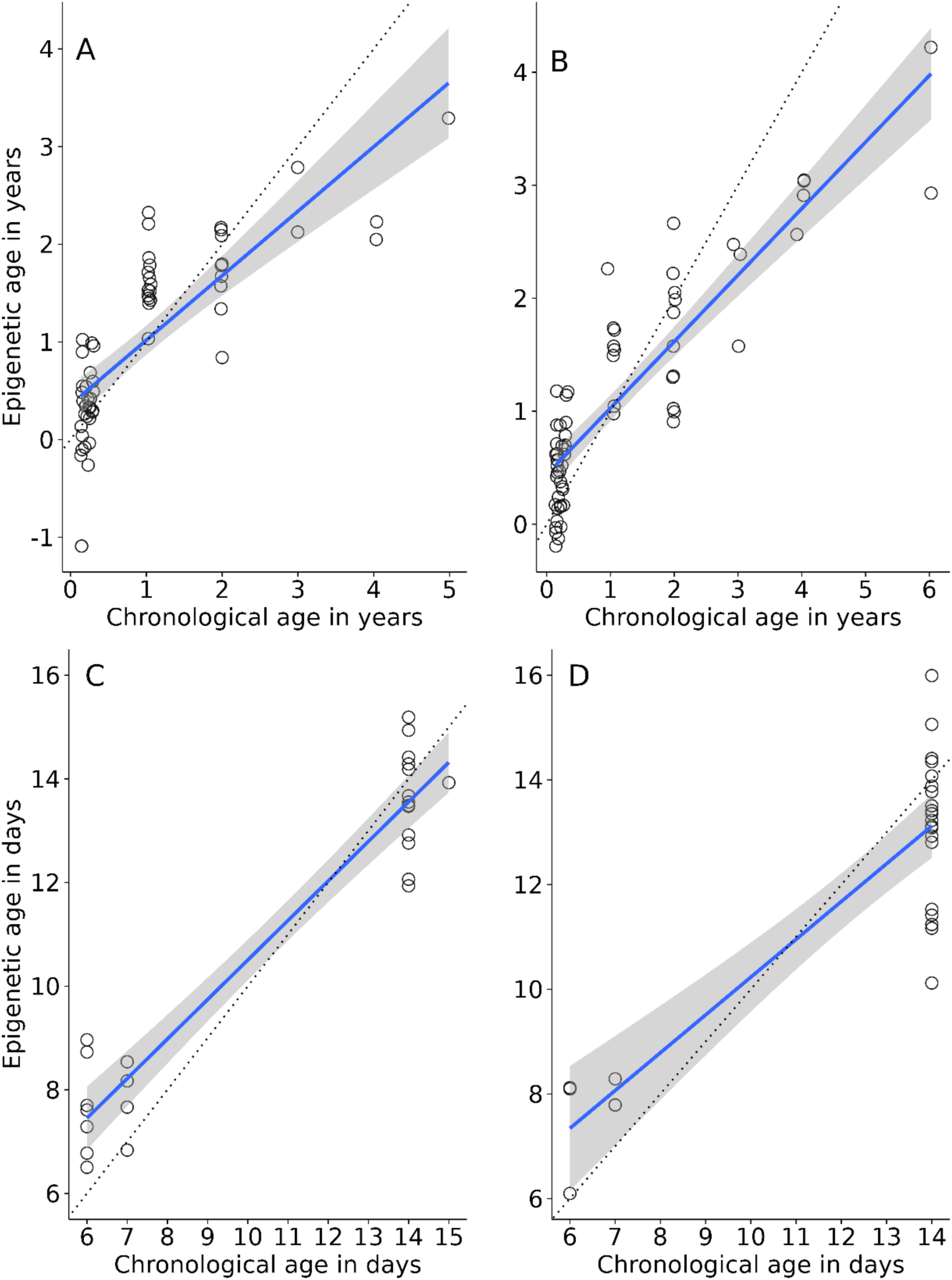
Epigenetic ages estimated using LOOCV for post-fledgling females (A; N = 58) and males (B; N = 64), and for pre-fledgling females (C; N = 25) and males (D; N = 26). Regression line of epigenetic age against chronological age is shown in blue with the standard error in grey. The dotted line represents the regression line when the epigenetic age is identical to the chronological age.

### 3.3 Testing the developmental clock

The 115 CpGs that were retained in the final development clock model were present in 96 individuals that were sampled at 14-days old, as part of the brood size manipulation experiment. Their epigenetic ages were estimated based on the determined coefficients of the final development clock. We found no significant difference in the mean epigenetic age between control and enlarged treatments (estimate = -0.254, *p* = 0.38), and between enlarged and reduced treatments (estimate = -0.33, *p* = 0.1). However, the reduced treatment showed to have a significantly higher epigenetic age compared to the control treatment (estimate = -0.58, *p* = 0.047) (Figure 4; Table 1).

**Figure 4.**
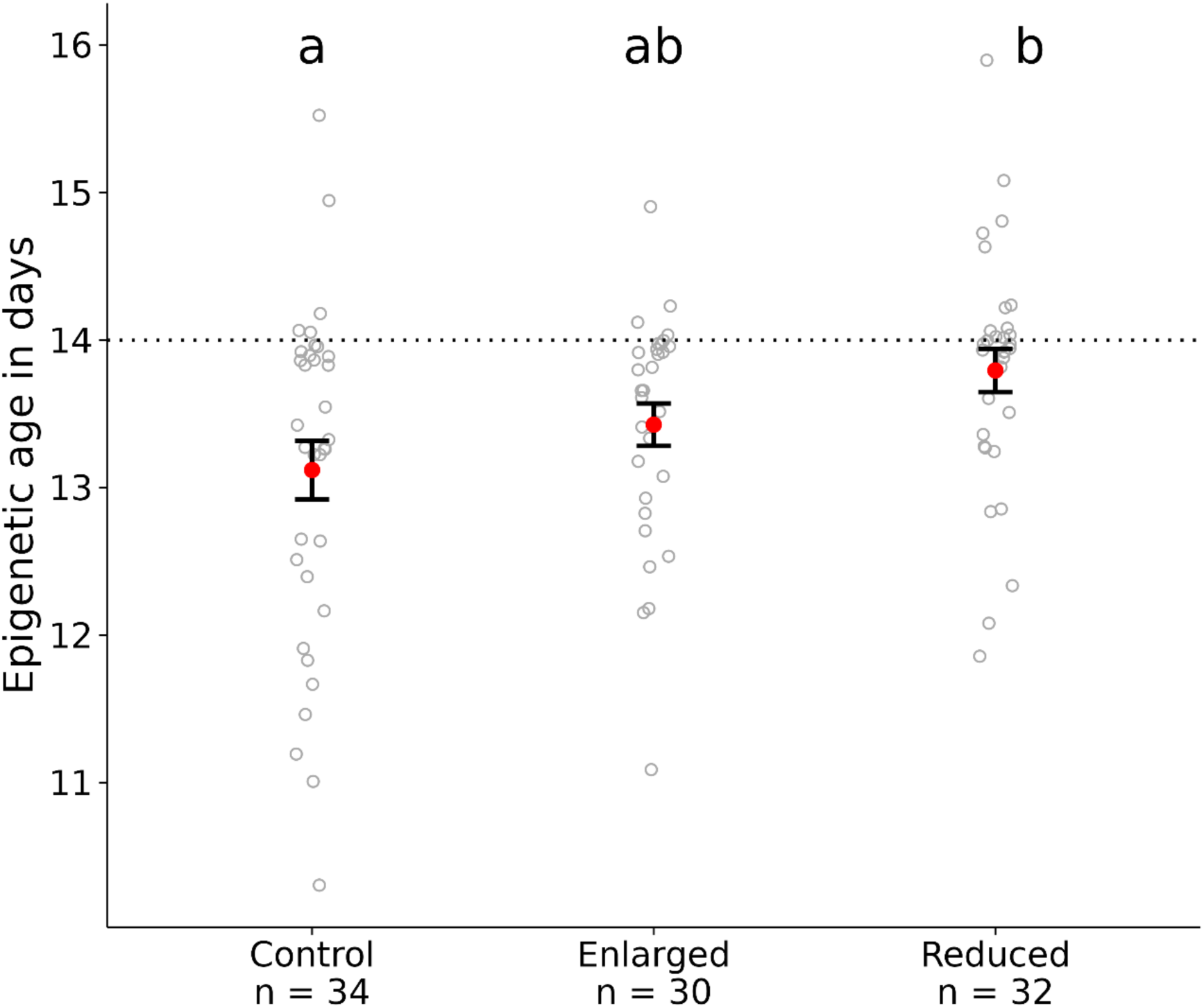
Mean (± standard error) epigenetic ages (shown in red), estimated using the final pre-fledgeling clock model, of 14-day old individuals in the control, enlarged and reduced treatment groups of a brood size manipulation experiment. Treatments with different letters are significantly different from each other, tested using the estimated marginal means with Fisher’s least significant difference test. Grey dots represent the raw estimated epigenetic ages of each individual.

**Table 1.** On the left a summary of the linear mixed model regressing epigenetic age against the different treatment groups, with brood of origin as random intercept effect, fitted using restricted maximum likelihood with a Gaussian error distribution. On the right the results of the post-hoc test, comparing the estimated marginal means (emmeans) with the lower and upper confidence intervals (CI-low and CI-up) between the control (N = 34), enlarged (N = 30) and reduced (N = 32) treatment groups. Treatments with a different letter under Sign. are significantly different from each other, tested with Fisher’s least significant difference test.

| Linear mixed model |  |  |  | Post-hoc test |  |  |  |  |  |
| --- | --- | --- | --- | --- | --- | --- | --- | --- | --- |
| Fixed Effects | Estimate | SE | t | Treatment | emmean | SE | CI-low | CI-up | Sign. |
| Intercept | 13.19 | 0.2 | 63.25 | Control | 13.2 | 0.21 | 12.8 | 13.6 | A |
| Enlarged | 0.25 | 0.3 | 0.89 | Enlarged | 13.5 | 0.20 | 13.1 | 13.8 | AB |
| Reduced | 0.58 | 0.3 | 2.05 | Reduced | 13.8 | 0.19 | 13.4 | 14.2 | B |
| Random Effect | $\sigma^2$ | $\sigma$ | $N$ | Anova Table | | | | | |
| Brood of origin | 0.36 | 0.6 | 34 |  | df | F-Statistic | P-value |  |  |
| Residuals | 0.56 | 0.8 |  | Treatment | 2 | 2.644 | 0.076 |  |  |
|  |  |  |  | Residuals | 94 |  |  |  |  |
| Linear mixed model |  |  |  | Post-hoc test |  |  |  |  |  |
| Fixed Effects | Estimate | SE | t | Treatment | emmean | SE | CI-low | CI-up | Sign. |
| Intercept | 13.19 | 0.2 | 63.25 | Control | 13.2 | 0.21 | 12.8 | 13.6 | A |
| Enlarged | 0.25 | 0.3 | 0.89 | Enlarged | 13.5 | 0.20 | 13.1 | 13.8 | AB |
| Reduced | 0.58 | 0.3 | 2.05 | Reduced | 13.8 | 0.19 | 13.4 | 14.2 | B |
| Random Effect | $\sigma^2$ | $\sigma$ | $N$ | Anova Table | | | | | |
| Brood of origin | 0.36 | 0.6 | 34 |  | df | F-Statistic | P-value |  |  |
| Residuals | 0.56 | 0.8 |  | Treatment | 2 | 2.644 | 0.076 |  |  |
|  |  |  |  | Residuals | 94 |  |  |  |  |

## 4. Discussion

The majority of research on the development of epigenetic clocks has been conducted on mammalian species (Lu et al., 2013). More recently, there has been a growing interest in the development of these clocks for other species as well. Consequently, epigenetic clocks for wild vertebrates such as fish (Mayne et al., 2023), a reptile (Mayne et al., 2022), and notably for two seabird species (De Paoli-Iseppi et al., 2019; Roman et al., 2024), have already been developed. In this study we used blood samples from pre- and post-fledging great tits (*P. major*) from a wild population to develop two distinct epigenetic clocks. These clocks provide further evidence that, as observed in mammals, DNAm changes of certain CpGs are highly correlated with chronological age in birds. Furthermore, we observed that individuals reared in experimentally reduced brood sizes exhibited higher epigenetic ages compared to individuals from control broods, but not compared to individuals from enlarged broods.

### 4.1 Epigenetic ageing models

To our knowledge, this is the first study to differentiate between an epigenetic ageing clock, based on samples from full grown individuals, and an epigenetic developmental clock, based on samples originating from developing individuals, within a wild species. The LOOCV approach allowed us to accurately predict the ages of pre- and post- fledglings (*r* = 0.89 and *r* = 0.84, respectively) with a MAD of 1.1 days and 0.4 years, respectively. The observed correlations were similar to those reported in clock models developed for mammals, such as Indo-Pacific bottlenose dolphins (*r* = 0.86; Peters et al., 2022) and beluga whales (*r* = 0.86; Bors et al. 2021). However, our clocks marginally outperform the first epigenetic clock that was constructed for a wild bird, short-tailed shearwater (*Ardenna tenuirostris*), that reported a coefficient of determination of 0.59 (De Paoli-Iseppi et al., 2019). Considering that the collective is greater than the contribution of individual parts (Horvath & Raj, 2018), it is generally thought that the accuracy of epigenetic clocks increases when more CpGs are included (Han et al., 2020; Horvath, 2013). In comparison to the *A. tenuirostris* clock, which only used seven CpGs in the final clock model, both our clock models included considerably more CpGs (183 CpGs in the ageing clock and 115 CpGs in the developmental clock), which may have accounted for the higher correlation. However, there is a trade-off between the accuracy and applicability between the number of CpGs that are included in the epigenetic clock models. Since a larger number of CpGs might increase the overall accuracy, while a smaller subset of CpGs may be easier to identify in additional samples, especially when a stochastic sequencing method, such as epiGBS2, is used to process the samples.

Despite the shorter lifespan of great tits compared to other species commonly used to construct epigenetic clocks, the error margins associated with the predictions (MAD) were relatively similar to those of other epigenetic clock models. Compared to the chronological ages, however, both clock models generally overestimated the age of younger individuals and the ageing clock underestimated the age of older individuals. This indicates that, in great tits, the epigenetic ageing does not follow a linear trend across different stages of the lifespan, and that the relationship is steeper in still-developing juveniles compared to fully-grown adults. The same changing rate of epigenetic ageing across different ages can be observed in other clock models, even in humans, where the ticking rate of the clock is higher during the development phase compared to adulthood (Horvath, 2013), possibly due to a high rate of mitotic divisions and the need for resource allocation during development (Lemaître, et al., 2022). On the other hand, the common underestimation of age in older individuals could reflect a survivor bias, where the lower epigenetic age of older individuals allowed them to live longer (Simpson & Chandra, 2021).

In our analysis we did not specifically exclude CpGs that were known to be located on a sex chromosome. However, none of the retained CpGs from both the final clock models were found to be located on a sex chromosome. Furthermore, our findings did not indicate a difference in the trend of the epigenetic ageing process between the estimated ages of male and female individuals. Evidence suggests that in certain species the process of epigenetic ageing is not equivalent in males and females (Horvath et al., 2016; Beal et al., 2019), and it is evident that in mammalian species females generally live longer than males (Lemaître et al., 2020). Moreover, it has been proposed that this may be dependent on the disparities in energetic expenditure between the sexes (Anderson et al., 2021). However, there have been few reports of longevity differences between sexes in birds (Austad & Fischer, 2016), and it remains unclear how longevity is related to genetic and epigenetic ageing processes. Meyer et al. (2023), demonstrated that there was a sex-specific difference in the global autosomal methylation rate in ageing common terns (*Sterna hirundo*), but reported no difference in senescence, nor average lifespan, in their study population. The development of these epigenetic clocks utilised cross-sectional samples and analytic tools that do not distinguish between differences in methylation patterns between sexes. Consequently, further research would be required to investigate sex-related disparities in the ageing process of great tits, and whether this can be related to the differences of sex-specific roles.

One limitation in the development of our epigenetic clocks was the unavailability of an equal distribution of ages in our samples. In case of the ageing clock model, there were abundant samples from younger individuals, but the number of samples from older individuals were limited. Whereas for the development clock model, we had a discontinuous distribution of ages in the samples. It has been demonstrated that the accuracy of epigenetic clocks improves when a broad range of ages, with an even distribution among the different age classes, are incorporated in the modelling data (Zhang et al., 2019). Nevertheless, the youngest and oldest individuals are generally prone to less accurate age estimations (Horvath, 2013), as was also the case in our models. An even distribution across the range of age classes that are included can increase the clocks correlation with chronological age (Lemaître et al., 2022), since the standard deviation of the age distribution is directly related to the clock accuracy (Tangili et al., 2023). However, as is the case with the majority of natural populations of great tits that have been studied, blood samples are mostly taken from nestlings during certain days after hatching, and from breeding adults. This results in a natural imbalance in the number of available samples throughout the whole developmental stage of nestlings, and there are often a greater number of samples taken from young individuals than from older ones. Such an imbalance may limit our understanding of the true epigenetic ageing processes throughout the whole nestling development, as well as our understanding of the ageing process in older individuals. Increasing our efforts to sample nestlings of different ages, and to track and sample older individuals in the population is therefore paramount to strengthen our understanding of the epigenetic ageing process throughout the whole lifespan of great tits.

### 4.2 Experimental validation of the epigenetic development clock

To validate the developmental clock, we quantified the epigenetic age of nestlings that were part of a brood size manipulation experiment (Sepers et al., 2023a), and assessed whether experimentally induced nutritional stress (Sepers et al., 2023b) influenced the epigenetic development process in early-life individuals. Nestlings reared in a reduced brood size had, on average, higher epigenetic ages compared to nestlings reared in an enlarged and control brood. However, this difference was only significant between the reduced and control treatments. Studies in humans suggest that stress is associated with increased epigenetic ageing (Zannas et al., 2015), but in the case of these nestlings, the individuals from a less stressful environments appeared to have the highest epigenetic age. We hypothesize that this reflects a faster development, which is expected to correlate with higher fitness and potentially lower ageing rates later in life. It is evident that epigenetic ageing is more dynamic during the developmental stage, however, further research is required to identify the association between environmental factors and the process of epigenetic ageing and development, as well as the impact of this process on the timing of key life history events and fitness.

## 5. Conclusion

This study demonstrates that the epigenetic age of great tits can be estimated based on DNAm, resulting in a high correlation with chronological age both before and after fledging. This is the first instance of an epigenetic clock being constructed for a wild passerine species. Given that various natural populations of great tits have been extensively studied and that there is abundant data for this model species, the clock model can be applied to analyse influences of epigenetic age on behavioural phenotypes, environmental factors and potential fitness consequences. Future improvements to these clocks could potentially reduce the number of highly correlated CpGs that are used, which would make the final clock models more applicable to samples that are sequenced with stochastic methods such as epiGBS2. Furthermore, it would be of interest to identify the specific genes that are the most closely associated with the ageing process to better understand the overall health and survival of individuals.

## Author contributions

KO, JR and AH designed research. BS collected the samples used for this study. AH and BS analyzed the data. AH conducted the statistical analysis. KO and JR supervised the study. AH wrote the first draft of the manuscript. All authors contributed to editing the manuscript.

## Acknowledgements

We gratefully acknowledge Nina Bircher, Piet de Goede and Youri van der Horst for fieldwork assistance. We thank Christa Mateman and Martijn van der Sluijs for their assistance in the lab and Rebecca Chen for helping with the bioinformatics analysis. We would like to thank Geldersch Landschap & Kasteelen for the permission to conduct fieldwork in Westerheide. This research was supported by an NWO-ALW open competition grant (ALWOP.314) to KvO.

## Data availability statement

Multiplexed reads are available on NCBI under BioProject ID PRJNA208335, under accession numbers SRX18523280, SRX18523281, SRX18523282, SRX18523283, SRX18523284, SRX18523285, SRX18523286, SRX18523287, SRX21758799, SRX21758800, SRX22027777, SRX22027778, SRX22027779, SRX22027780, SRX22027781, SRX26112229, SRX26112230, SRX26112231, SRX26112232, SRX26112233 and SRX26112234. Barcode files, R scripts and sample datasheets used to analyse the methylation calls and to construct the epigenetic clocks will be made available upon publication. The epiGBS2 pipeline can be accessed on Github (https://github.com/nioo-knaw/epiGBS2).

## Ethics statement

All animal experiments involved in this study were reviewed and approved by the Institutional Animal Care and Use Committee (NIOO-IvD) and were licenced by the CCD (Central Authority for Scientific Procedures on Animals; AVD-801002017831) to KO.

