## Supplementary materials for "Independent avian epigenetic clocks for aging and development"

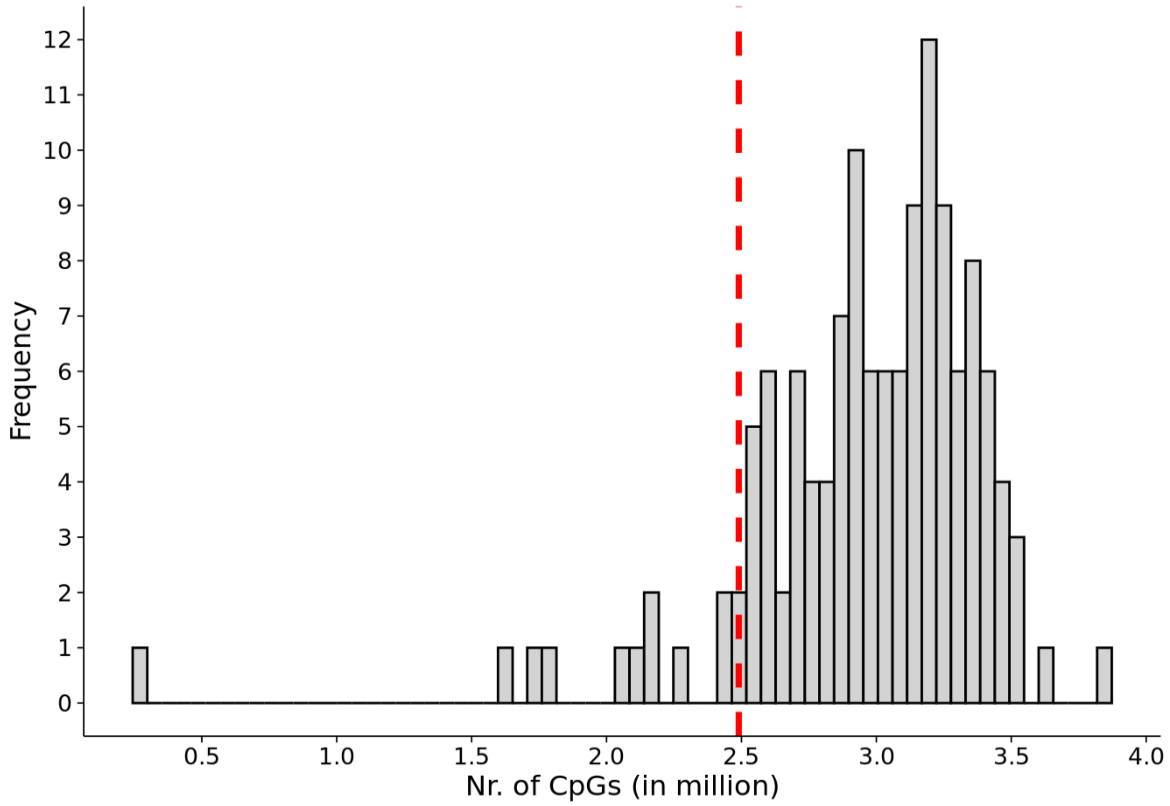


Figure S1. Histogram of the number of unmerged CpGs from 134 post-fledging samples. The red line indicates the cut-off point of 12 samples that were removed from the analysis.

Table S1. The number of CpGs of the post-fledging datafiles and the number of remaining CpGs after each filtering step. Coverage filter, high percentile of 99.9% filter, filter for mean methylation more than 5% and less than 95%, and filter for standard deviation of more than 0.05, were only applied to the CpGs that were covered throughout the remaining 122 samples.

| | **Average** | ***N* samples** |
|---|---|---|
| Total CpGs | 15,372,018 | 134 |
| Empty CpGs removed | 2,963,751 | 134 |
| Merged CpGs | 2,552,100 | 122 |
| **Selected CpGs only in all samples** | | |
| CpGs (10x min. coverage & high percentile filter of 99.9%) | 35,126 | 122 |
| Mean & SD filter | 8,398 | 122 |

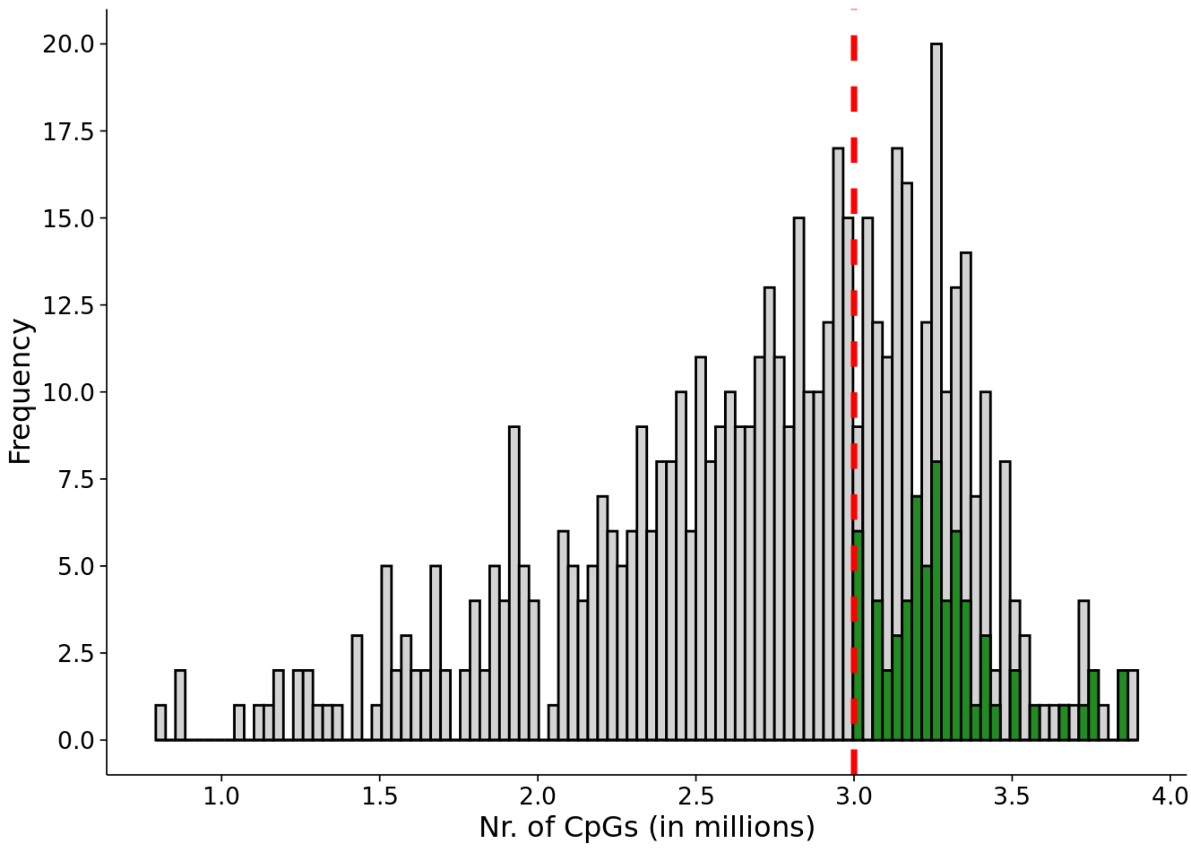


Figure S2. Histogram of the number of unmerged CpGs from 618 pre-fledging samples. The red line indicates the cut-off point of 346 samples that were excluded from the analysis. Green bars indicate the 67 samples that were selected to construct the clock model.

Table S2. The number of CpGs of pre-fledging datafiles and the number of remaining CpGs after removing empty CpGs with 0 Cs and Ts, and the number of CpGs that were present in at least one sample after filtering for >10 coverage and a coverage over high percentile of 99.9%.

| | **Average** | ***N* samples** |
|---|---|---|
| Total CpGs | 15,372,018 | 618 |
| Empty s removed | 2,784,984 | 618 |
| **Selected samples > 3 million CpGs** | | |
| Merged & Filtered CpGs (10x min. coverage & high percentile filter of 99.9%) | 2,055,751 | 272 |

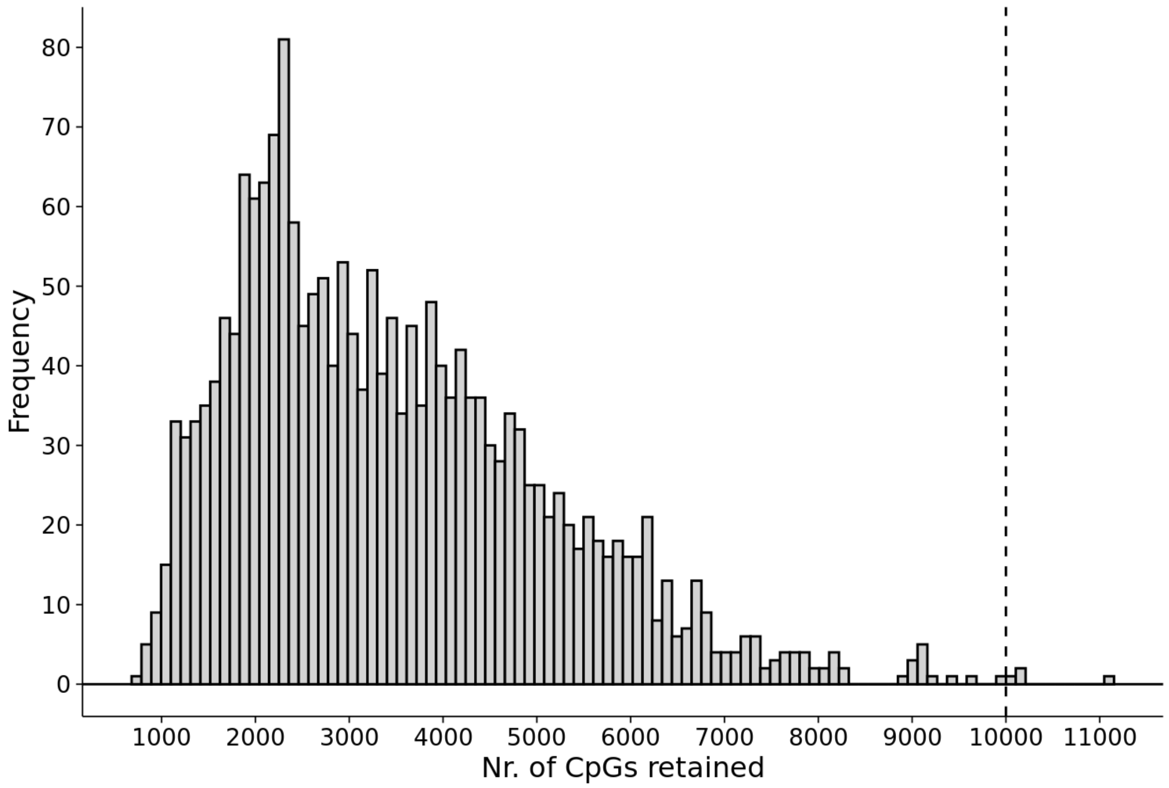


Figure S3. Histogram of numbers of CpGs that were retained in 500 randomised subsets of ~25% from 272 pre-fledglings. Four subsets were found that contained > 10,000 CpGs, indicated by the dashed lined.

Table S3. The number CpGs in the selected subset of pre-fledging individuals that were used to construct the final clock model.

| | **Number** | ***N* samples** |
|---|---|---|
| Total CpGs | 11,116 | 67 |
| Filtered CpGs (10x min. coverage & high percentile filter of 99.9%) | 3,184 | 67 |

Table S4. Number of pre-fledging samples per experiment that were used to construct the epigenetic clock.

| | ***N*** | **Age range in days** | **Males** | **Females** | **Unknown** |
|---|---|---|---|---|---|
| All nestling samples | 67 | 6 - 15 | 26 | 25 | 16 |
| **Per Experiment** | | | | | |
| Brood size manipulation | 34 | 14 – 15 | 20 | 14 | |
| Testosterone manipulation | 17 | 6 – 7 | 6 | 11 | |
| Food deprivation | 16 | 6 - 7 | | | 16 |
